# Microbiomes in the insectivorous bat species *Mops condylurus* rapidly converge in captivity

**DOI:** 10.1101/784074

**Authors:** Kathryn M Edenborough, Andre Mu, Kristin Mühldorfer, Johanna Lechner, Angelika Lander, Marcel Bokelmann, Emmanuel Couacy-Hymann, Aleksandar Radonic, Andreas Kurth

## Abstract

Bats are well known reservoir hosts for RNA and DNA viruses. The use of captive bats in research has intensified over the past decade as researchers aim to examine the virus-reservoir host interface. In this study, we investigated the effects of captivity on the fecal bacterial microbiome of an insectivorous microbat, *Mops condylurus*, a bat species that roosts in close proximity to humans and has likely transmitted viral infections to humans. Using amplicon 16S rRNA gene sequencing, we characterized changes in fecal bacterial community composition for individual bats directly at the time of capture and again after six weeks in captivity. We found that microbial community richness by measure of the number of observed operational taxonomic units (OTUs) in bat feces does increase significantly in captivity. Importantly, we found the similarity of microbial community structures of fecal microbiomes between different bats to converge during captivity. We propose a six week-acclimatization period prior to carrying out infection studies or other research influenced by the microbiome composition, which may be advantageous to reduce variation in microbiome composition and minimize biological variation inherent to *in vivo* experimental studies.

## Introduction

Bats (*Mammalia, Chiroptera*) play an important role in pollination and pest control (1) and are natural host reservoirs for many RNA and DNA viruses known to cause significant morbidity and mortality in humans (2). To effectively study their role as reservoir hosts for pathogenic viruses, bats have been housed in captivity (3–6). Introduction of bats to captive environments entails dietary adaptation, increased stress and habitat change (7, 8). The captive environment also indirectly modifies the microbiome with respect to the number of microbial species in the community (alpha diversity) and the phylogenetic similarity of the microbial communities between separate bats (beta diversity) (9, 10).

The fecal microbiomes in individual wild bats are unique and cluster based on host phylogeny and feeding strategy (11), however in captivity the microbial communities have been shown to converge over six months (9). Convergence to shared microbial communities by the captive group is largely influenced by diet (based on phylogenetically aware diversity metrics such as UniFrac). For captive studies, bats are typically introduced into captivity over time and therefore the individuals originate from different geographical locations and are likely exposed to regional diets. Identifying a minimum period of housing for the microbiome to converge in captivity would assist researchers to choose more accurate acclimatization periods to prevent the influence of interanimal variation of the microbiome from confounding experimental findings.

Our current knowledge of microbial convergence of the fecal microbiome in captive bats extends to a six-month window and we therefore sought to measure changes to the fecal microbiome in captivity within a shorter time frame. We also focused on the bat species *Mops condylurus (M. condylurus*) that roosts in close proximity to villages and has likely transmitted viral infections to humans in the past (12). Fecal samples were collected from insectivorous bats prior to capture (pre-capture samples) and following six weeks in captivity (post-capture samples) and using amplicon 16S rRNA gene sequencing to interrogate the microbiota, we found the microbial communities in bats became phylogenetically similar within six weeks of captivity.

## Materials and Methods

### Sample collection

Animal capture, handling and sampling were performed with the permission of the Laboratoire Central Vétérinair, Laboratoire National D’Appui Au Developpement Agrcole (LANADA), Bingerville, Ivory Coast according to ethics application (No. 05/Virology/2016). The animal care and use protocol adhered with the ethics committee of LANADA. *Mops condylurus* bats were captured with mist nets near Koffikro, Ivory Coast, and held individually in cotton bags until transfer to captivity at the Le Laboratoire Central Vétérinaire de Bingerville (LCVB). Bats were held in a steel framed meshed aviary, sized 5 metres in length and 4 metres in width. Roosting boxes and hanging soft material were provided as hiding places for the bats. The captive diet consisted of mealworms (*Tenebrio molitor*) which were fed to bats with tweezers or were first made into a mealworm puree and fed to bats via syringe until the point of satisfaction. Fecal samples were collected twice; directly after capture (pre-capture) and at six weeks of captivity (post-capture) for microbial community profiling using amplicon 16S rRNA gene sequencing. Samples were obtained by collection of fecal pellets (~100-300 mg) in screw cap 2 ml micro tubes (Sarstedt) from cotton bags used to hold individual microbats for about two hours. After collection fecal samples were frozen at −80°C and later transported to the Robert Koch Institute, Berlin in a cryogenic dry shipper where they were stored at −80°C until genomic DNA extraction.

### Metadata for microbiome datasets

Fecal pellets were collected directly from 20 individual *M. condylurus* bats upon capture (herein referred to as pre-capture samples) and from the same animal six weeks after introduction to captivity (herein referred to as post-capture samples) following adaptation to the feeding program. There was sufficient fecal material to perform nucleic acid extractions for nine pre-capture and 18 post-capture bats. All bats were species confirmed as *M. condylurus* by amplification and sequencing of the cytochrome b gene. The metadata for these samples is shown in **Table S1**.

### Nucleic acid extraction

Genomic DNA was extracted from ~100 mg of feces with the use of a Nucleospin DNA Stool Kit (740472, Macherey-Nagel) according to the manufacturer’s instructions with the following modifications; for sample preparation each Nucleospin Tube Type A containing 100 mg of feces was loaded into pre-chilled adapters and shaken twice for 30 sec with a TissueLyser II instrument (Qiagen) at a frequency of 25 Hz. Following lysis at 70°C for 5 mins, the samples were further disrupted by vigorous shaking with a ThermoMixer (Eppendorf) at room-temperature, 2000 rpm for 10 min. Nucleic acids were eluted in 50 μl elution buffer and stored at −20°C.

### Small Subunit 16S rRNA gene V3-V4 amplification with fusion primers

Amplification and purification of V3-V4 regions of the 16S rRNA genes were performed according to the 16S Metagenomic Sequencing Library Preparation protocol (13) from Illumina ^®^ with the following modifications; PCR was carried out with Kapa High Fidelity (HiFi) PCR Kit (KK2101, Kapa Biosystems) by the addition of 25 ng DNA to a master mix containing 1×HiFi Buffer, 2.25 mM magnesium chloride, 0.3 μM of each forward and reverse primer, 1.2 mM deoxynucleotides and 0.5U KAPA HiFi DNA Polymerase. PCR conditions included one cycle of 95°C for 3 min, 30 cycles of 98°C 30 sec, 57°C 30 sec, 72°C 30 sec, and one cycle of 72°C for 5 min. Forward 5’-TCGTCGGCAGCGTCAGATGTGTATAAGAGACAGCCT ACGGGNGGCWGCAG-3’ and reverse 5’-GTCTCGTGGGCTCGGAGATGTGTATAA GAGACAGGACTACHVGGGTATCTAATCC-3’ primers were based on previously optimized sequences (14).

### Library preparation

Amplicons were subjected to index PCR, library pooling according to Illumina protocols (13) and paired-end sequencing on a MiSeq instrument with MiSeq Reagent Kit v3 (MS-102-3003, Illumina). The indices for each sample are provided in **Table S1**

### Data availability

The fastq files for amplicon 16S rRNA gene sequencing are available in the sequence read archive repository, accessible under Bioproject PRJNA516511.

### Microbiota analyses

Bioinformatic analyses of 16S rRNA gene data were carried out on demultiplexed fastq reads. Reads were processed with Skewer version 0.2.1 to filter out reads (Q < 12) and to remove adapter and primer sequences. Reads were analyzed with QIIME2 (version 2019.1.0) by implementing plug-ins including Deblur to denoise sequencing reads (15), SEPP to phylogenetically place reads (16), feature-classifier to taxonomically assign OTUs, UniFrac to compute beta-diversity analyses (17) and gneiss and balance trees (18) to examine relative abundances between variables. Further statistical tests were carried out in QIIME2 including Kruskal-Wallis pairwise test for Faith’s PD, pairwise PERMANOVA to compare beta-diversity and Prism (version 8.8.0) for comparison of sequence reads with Mann-Whitney test.

## Results

### Fecal microbiomes of M. condylurus in captivity have greater species richness than in the wild

The number of 16S rRNA gene reads sequenced for each bat fecal sample in pre-capture (45×10^3^ ± 6.7×10^3^) and post-capture (54×10^3^ ± 28×10^3^) groups are shown in **Fig 1A**. The mean number of reads sequenced for pre-capture and post-capture groups are comparable (not significantly different Mann-Whitney test), although variance in sequence read numbers for the post-capture group is high, thus the data is subsampled at a depth of 4000 sequences to standardize read number between the two groups. A subsampling depth of 4000 reads recovers the majority of OTUs for each sample as shown by transition to a plateau in the rarefaction curves presented in **Fig 1B** and this subsampling depth has been implemented to calculate various diversity metrics.

**Fig 1.**
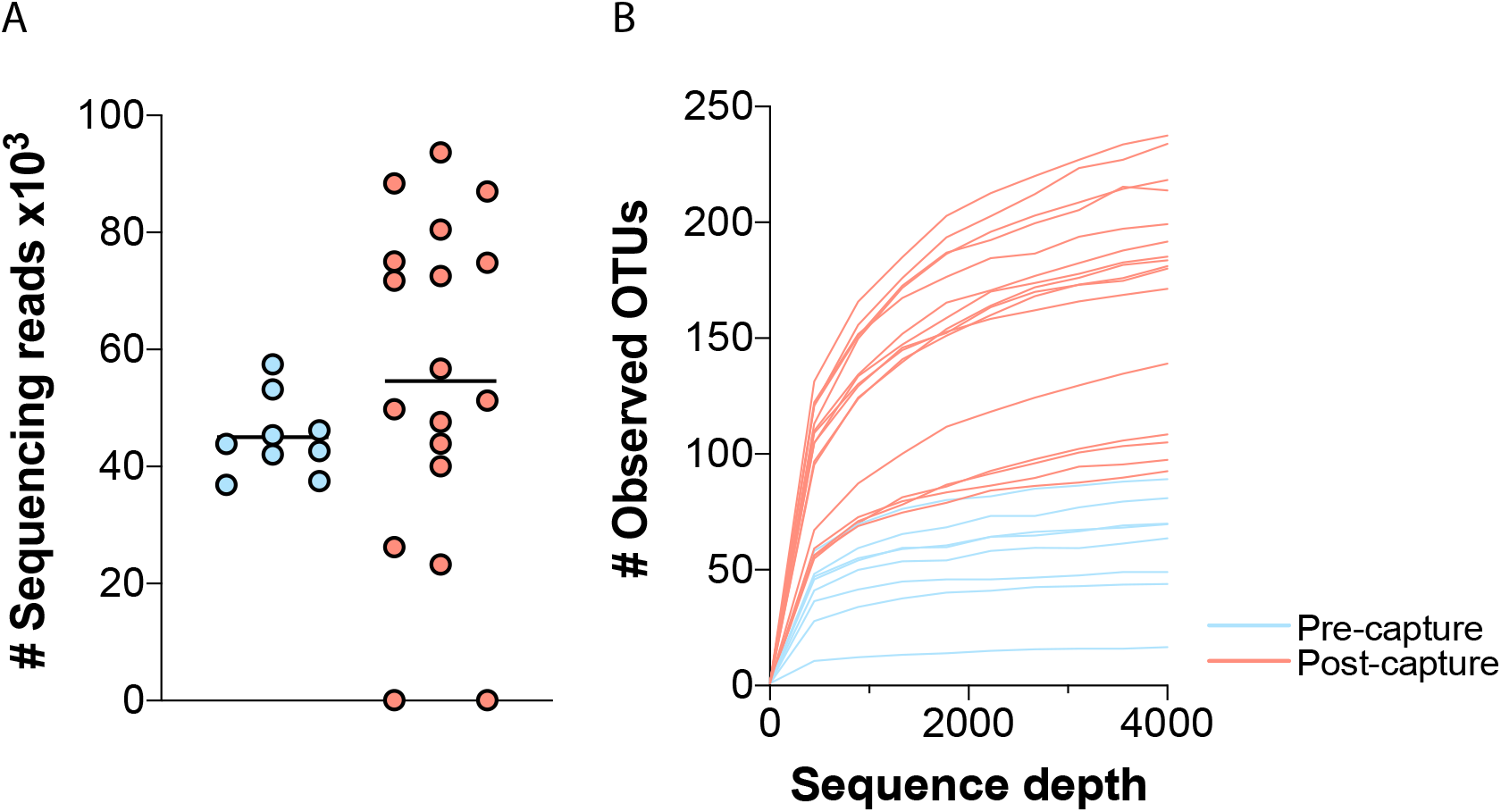
Number of sequencing reads and mapped OTUs in fecal bat samples. In **A** number of 16S rRNA V3-V4 gene reads sequenced for bat fecal samples collected prior to capture (light-blue) and post-capture (light-red) are shown for individual bats in a scatter plot. In **B** the number of observed OTUs recovered from data rarefied at a sequence depth of up to 4000 sequences are shown for each pre-capture (light-blue) and post-capture (light-red) sample.

Alpha diversity metrics are calculated with QIIME2 to compare microbial diversity for individual bats within their pre-capture and post-capture microbiotas. Shown in **Fig 2A** are Faith’s phylogenetic diversity (PD) scores, which enumerate microbial richness by considering branch length and relatedness of the OTUs within the phylogenetic tree. Faith’s PD values for post-capture microbiomes are significantly higher, indicating increased species richness than pre-capture microbiomes (p = 0.000089, ANOVA). No significant differences in Faith’s PD values exist for other variables such as gender, mealworm consumption, bodyweight and time of capture (data not shown).

**Fig 2.**
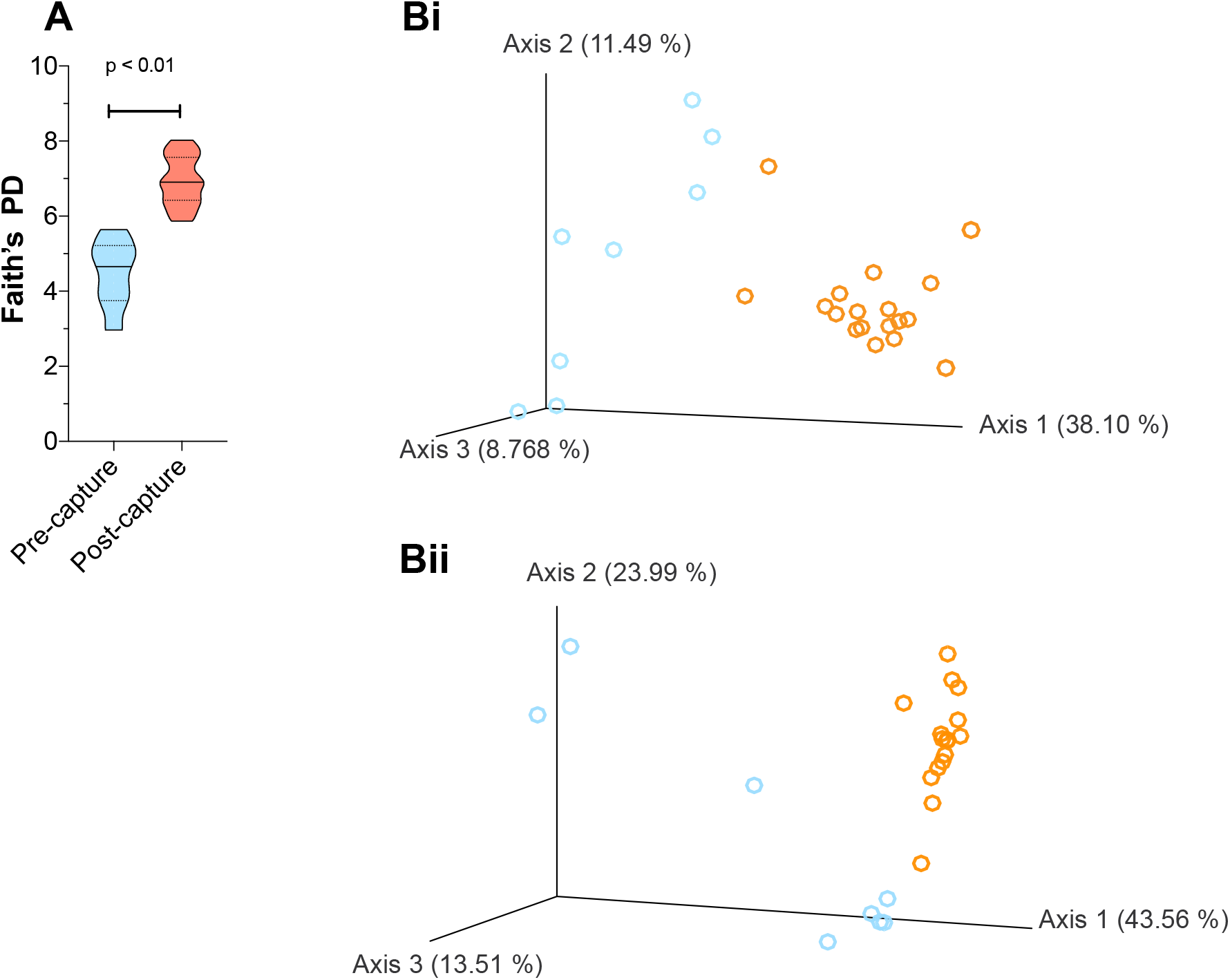
Alpha and beta diversity metrics for pre-capture and post-capture datasets. With a violin plot **(A)** Faith’s phylogenetic diversity (PD) score is shown for OTU mapped 16S rRNA gene sequences amplified from fecal samples, which were collected from nine pre-capture bats and 18 post-capture bats. The beta diversity of microbial communities in each pre-capture (light-blue) and post-capture (light-red) sample was calculated with unweighted UniFrac (**Bi**) and weighted UniFrac (**Bii**) and scaled values are shown with a PCoA plot visualized with Qiime2view.

### Convergence of fecal microbiomes for microbats housed in captivity

Beta diversity metrics are used to measure the dissimilarity of microbial communities in individual animals. Beta diversity is calculated with QIIME2 by implementing unweighted- and weighted-UniFrac analyses and the results are shown in **Fig 2Bi and 2Bii**. Unweighted UniFrac data scaled with principal coordinates analysis (PCoA) reveals the first factor accounts for 38.1% of variation in the data (**Fig 2Bi, axis 1**). Post-capture samples form a distinct cluster on axis 1, while precapture samples disperse across the first and second axes. The difference observed between precapture and post-capture microbial communities on scaled PCoA plots is further supported by a significant difference between the two groups in UniFrac distance metrics (p<0.001, PERMANOVA). Clustering of post-capture samples is also visible for analyses giving weight to microbial relatedness and microbial abundance, such as weighted UniFrac. In summary, these data reveal that post-capture, microbial species are significantly similar in taxonomic quality and abundance while at pre-capture the microbial communities are qualitatively diverse.

### Compositional changes associated with M. condylurus microbiome in captivity

UniFrac analyses revealed clustering of post-capture samples and we investigated the taxa contributing to this effect by examining the relative frequency of bacteria in pre-capture and postcapture samples. Relative frequencies of features identified at the level of bacterial class are shown for each pre-capture and post-capture sample in **Figure 3A**. The dominant bacterial phyla identified in all samples, regardless of capture status includes; Proteobacteria, Firmicutes and Actinobacteria. First-order level analyses of biodiversity profiling reveals that Actinobacteria are abundant in post-capture samples, while a specific class within the Firmicutes phyla, Clostridia are abundant in pre-capture samples.

**Fig 3.**
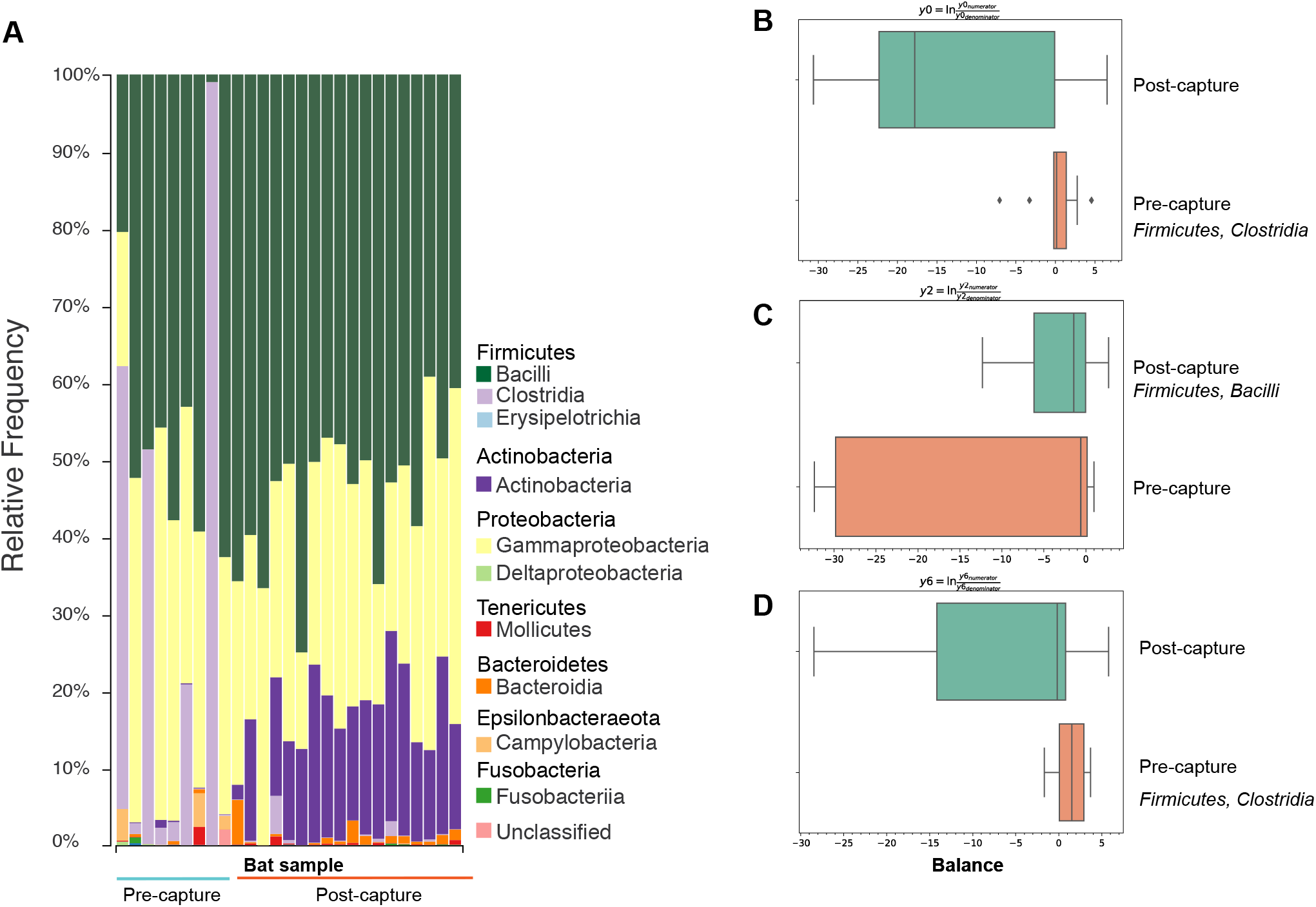
Taxonomical variation in abundance between fecal samples. The relative frequency of reads that mapped to classified features are shown in **A** at the class level for each bat sample collected pre-capture (light-blue horizontal bar) and post-capture (light-red horizontal bar). Feature abundance was clustered in reference to capture status and specific bacterial groups identified that were significantly different in balances y0 (B) y2 (C) and y6 (D) between pre-capture (orange) and post-capture (green) samples.

A deeper analysis of the microbial composition changes between pre-capture and post-capture states was performed with Gneiss (18). Initially a Ward-hierarchical clustered heatmap was produced, which shows the coefficient p-values for each sample in one column for each of the different balances y0-y9 (**Fig S1**). Balances y0, y2 and y6 are of particular interest and we therefore explored the taxa constituting these balances further in **Fig 3B-D**. The main taxon contributing to the microbial signature of pre-capture samples is Firmicutes, Clostridia which is observed at a greater proportion than in post-capture samples for both y0 and y6 balances. A key taxon revealing the microbial signature of the post-capture samples is Firmicutes, Bacilli, which is observed at greater proportions than in pre-capture samples for the y2 balance. *Actinobacteria*, Corynebacteriales and Actinomycetales were also identified in post-capture samples at slightly higher proportion than pre-capture samples. They likely culminate in the apparent increase in relative frequency in Actinobacteria within post-capture samples as seen in **Fig 3A**. Overall, interrogation of the microbial composition has revealed the fecal microbiota changes in captivity in relation to the wild microbial community. These changes are distinguished by a co-housing effect, where microbial profiles from individual animals converge to form an exceedingly homogenous bacterial community.

## Discussion

In this study, we investigated the influence of captivity on the fecal microbiota of microbats, *M. condylurus* by comparing microbial diversity in samples collected from bats on the day of capture and again from bats after six weeks of captivity. Using beta diversity metrics, we determined that the fecal microbial communities in pre-capture samples were phylogenetically diverse while those in post-capture samples were phylogenetically homogenous. For other mammals, such convergence of fecal microbial communities in captivity is largely explained by adaptation to a captive diet, residing in geographically close locations, and exposure to humans (10, 19). For bats, past studies have shown microbial composition is influenced by diet, environment, co-housing and feeding strategy (9, 20, 21). One particular study by Xiao et al shows that convergence of the microbial fecal population, for various microbat species co-housed in a laboratory environment is due to dietary change over a six-month period (9). One limitation of this study, and also our study here is that the microbial community of the ingested captive diet was not determined, and it is therefore unclear whether the fecal microbial species identified are commensal to the diet or are part of the native intestinal flora. Nevertheless, our study extends these findings by observing convergence during post-capture over a short time frame of six weeks. This has important implications for acclimatization time prior to embarking on experimental research with bats, which should be at least six weeks if the researcher intends to control for variations in microbial communities between different animals. On the other hand, if a wild microbiome is a prerequisite for a research outcome then performing research within six weeks after capture would be ideal. Despite the homogeneity observed between post-capture samples, these same samples were also identified as more species rich (higher number of OTUs detected) than pre-capture samples with alpha diversity metrics. This finding contrasts with the research investigating species richness of bacterial communities in terrestrial mammals, which has revealed animals found in the wild have greater microbial species richness than those kept in captivity and only in a small number of mammals (odd-toed ungulates) is stable with regards to its microbial community richness irrespective of captive status (10, 22). Rather, this finding echoes those presented in bat specific studies which revealed species richness to increase in captivity for fruit bats (20) and microbats (9, 23). These studies have suggested post-capture species richness is a result of shared microbiomes and commensal bacteria ingested as part of the captive diet. One alternative explanation is a greater range of bacterial species can expand in the absence of bacteria required to process a diet rich in chitin, which forms part of the exoskeleton of insects that are consumed by *M. condylurus* and most microbats. Notwithstanding, prior studies have demonstrated a reduction in bacterial species richness in captive fruit bats (24) and other studies suggest inconsistencies in alpha diversity metrics between studies are common and result from different analysis pipelines, whereas beta diversity metrics may be more comparable between studies (25). Microbial species classified within the Firmicutes and Proteobacteria phyla were commonly detected irrespective of capture status and this profile matches bat fecal microbiota profiles shown in other published works (9, 11, 21). The convergence of post-capture microbial landscapes was signified by increases in Bacilli and Actinobacteria spp. while the signature of pre-capture microbiomes was an abundance of Clostridia spp. The fluctuations in bacterial composition caused by captivity correspond to those seen in other captive mammals (10). The uniqueness to our dataset is the identification of a high frequency of Actinobacteria spp. in captive bats, which is more typically identified at a lower frequency. To understand the implications of finding Actinobacteria in abundance further interrogation into the commensal microbial communities of the captive diet would be required as insects are known to harbor an abundance of Proteobacteria spp. but not Actinobacteria spp. in their gut (26).

Our findings reveal that fecal microbial communities in microbats change and converge as a result of captivity. Taken together, these findings imply that with the use of a short co-housing period of six weeks the fecal microbiome will become similar between disparate bats co-housed in captivity. A six-week acclimatization period could be implemented to reduce the confounding effects of inter-animal variation in microbial communities for a more controlled experimental system.

## Acknowledgements

The authors would like to acknowledge Norman Kirchhoff, Nicole Hetzelt, Katharina Hansen-Kant, Romeo Yao and Assamoa for their persistence in bat catching and feeding of the captive colony and Brendon Chua for editing the manuscript. This research received no specific grant from any funding agency in the public, commercial, or not-for-profit sectors.

## Supporting Information Captions

**Fig S1.**
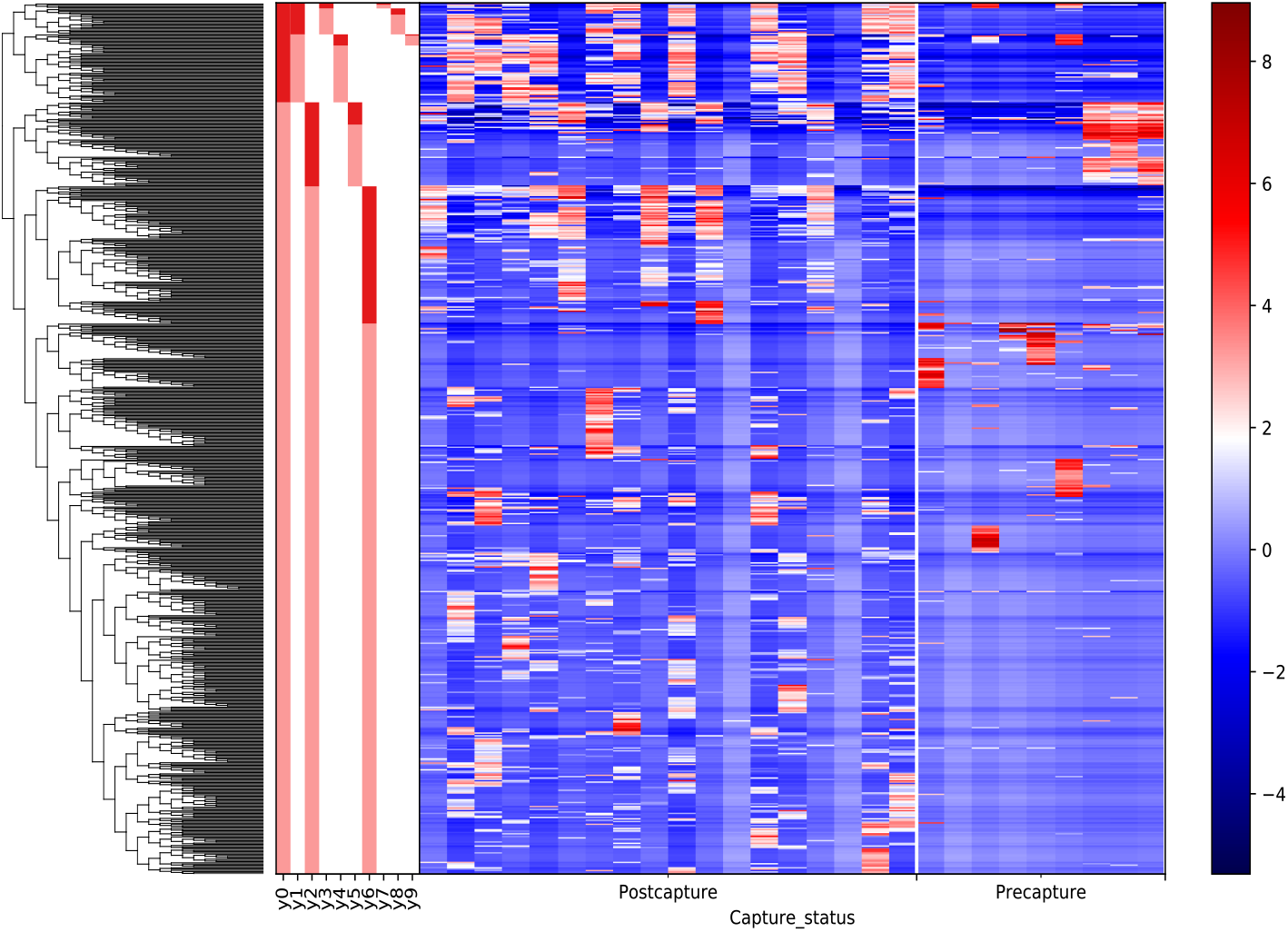
Gneiss heatmap of Ward clustered partitions. Using the gneiss toolkit in Qiime2 a heatmap was created and shows the coefficient p values for balances y0-y9 after comparison of pre-capture and post-capture samples. Each balance consists of clustered OTUs that strongly correlate in their abundance in pre-capture or post-capture samples.

## Declarations

### Ethics approval

Animal capture and procedures undertaken were permitted under authorization number 05/Virology/2016 granted by Laboratoire National D’Appui Au Developpement Agrcole (LANADA).

### Availability of data and materials

Supporting fastq files for the datasets generated in this article are available in the NCBI sequence read archive repository https://www.ncbi.nlm.nih.gov/sra, accessible under SRR8475248-263 belonging to Bioproject PRJNA516511.

### Competing Interests

The authors declare that they have no competing interests

### Funding

This publication was financially supported by the RKI

### Authors contributions

Conceptualization: AK, KM & KE, Data Curation and analysis: KE & AM, Funding: AK, Investigation: KE, AL, MB, AR and JL, Project administration: ECH & AK, Writing & editing: KE, AM & KM

---

**S1:** Associated metadata with 16Sr RNA sequence datasets.

S1: Associated metadata with 16Sr RNA sequence datasets
| Sample ID | Bat Number | Gender | Collection Date | Index1 | Index2 | Microbiome Status |
| --- | --- | --- | --- | --- | --- | --- |
| 1 | 1 | M | 7/4/17 | ACTCGCTA | CGTCTAAT | Pre-capture |
| 2 | 3 | F | 7/4/17 | ACTCGCTA | TCTCTCCG |  |
| 3 | 8 | F | 7/4/17 | GGAGCTAC | CTCTCTAT |  |
| 4 | 24 | M | 14/4/17 | GGAGCTAC | TATCCTCT |  |
| 5 | 25 | M | 14/4/17 | GGAGCTAC | GTAAGGAG |  |
| 6 | 26 | M | 14/4/17 | GGAGCTAC | ACTGCATA |  |
| 7 | 28 | M | 22/4/17 | GGAGCTAC | AAGGAGTA |  |
| 8 | 34 | M | 23/4/17 | GGAGCTAC | CTAAGCCT |  |
| 9 | 35 | F | 23/4/17 | GGAGCTAC | CGTCTAAT |  |
| 10 | 1 | M | 19/5/17 | GCGTAGTA | GTAAGGAG | Post-capture |
| 11 | 3 | F | 19/5/17 | GCGTAGTA | ACTGCATA |  |
| 12 | 8 | F | 19/5/17 | GCGTAGTA | AAGGAGTA |  |
| 13 | 13 | F | 19/5/17 | GCGTAGTA | CTAAGCCT |  |
| 14 | 14 | M | 19/5/17 | GCGTAGTA | CGTCTAAT |  |
| 15 | 15 | F | 19/5/17 | CGGAGCCT | CTCTCTAT |  |
| 16 | 18 | M | 19/5/17 | CGGAGCCT | TATCCTCT |  |
| 17 | 24 | M | 19/5/17 | CGGAGCCT | GTAAGGAG |  |
| 18 | 25 | M | 19/5/17 | CGGAGCCT | ACTGCATA |  |
| 19 | 26 | M | 19/5/17 | CGGAGCCT | AAGGAGTA |  |
| 20 | 28 | M | 19/5/17 | CGGAGCCT | CTAAGCCT |  |
| 21 | 33 | M | 19/5/17 | CGGAGCCT | CGTCTAAT |  |
| 22 | 34 | M | 19/5/17 | CGGAGCCT | TCTCTCCG |  |
| 23 | 35 | F | 19/5/17 | TACGCTGC | CTCTCTAT |  |
| 24 | 41 | M | 19/5/17 | TACGCTGC | TATCCTCT |  |
| 25 | 49 | M | 19/5/17 | TACGCTGC | GTAAGGAG |  |
| 26 | 50 | F | 19/5/17 | TACGCTGC | ACTGCATA |  |

## References

1. Kasso M, Balakrishnan M. Ecological and Economic Importance of Bats (Order Chiroptera). ISRN Biodivers. 2013; Article ID 187415. doi:10.1155/2013/187415

2. Wang L-F, Cowled C. Bats and Viruses: A New Frontier of Emerging Infectious Diseases. John Wiley and Sons Inc.; 2015. doi:10.1002/9781118818824

3. Schuh AJ, Amman BR, Jones MEB, Sealy TK, Uebelhoer LS, Spengler JR, et al. Modelling filovirus maintenance in nature by experimental transmission of Marburg virus between Egyptian rousette bats. Nat Commun. 2017; 8: 14446. doi:10.1038/ncomms14446

4. Jones EM, Schuh JA, Amman RB, Sealy KT, Zaki RS, Nichol TS, et al. Experimental Inoculation of Egyptian Rousette Bats (Rousettus aegyptiacus) with Viruses of the Ebolavirus and Marburgvirus Genera. Viruses. 2015; 7(7). doi:10.3390/v707277

5. Paweska TJ, Storm N, Grobbelaar AA, Markotter W, Kemp A, Jansen van Vuren P. Experimental Inoculation of Egyptian Fruit Bats (Rousettus aegyptiacus) with Ebola Virus. Viruses. 2016; 8(2). doi:10.3390/v8020029

6. Freuling C, Vos A, Johnson N, Kaipf I, Denzinger A, Neubert L, et al. Experimental infection of serotine bats (Eptesicus serotinus) with European bat lyssavirus type 1a. J Gen Virol. 2009; 90(10): 2493–502. doi:10.1099/vir.0.011510-0

7. Barnard SM. Bats in Captivity. Barnard S, editor. Logos Press; 2009. (Bats in Captivity).

8. Chen Y, Liu Q, Shao YG, Tan LJ, Xiang ZF, Zhang LB. Variation in echolocation calls of Hipposideros armiger during habituation to a novel, captive environment. Behaviour. 2015; 152(7–8): 1083–95. doi:10.1163/1568539X-00003269

9. Xiao Y, Xiao G, Liu H, Zhao X, Sun C, Tan X, et al. Captivity causes taxonomic and functional convergence of gut microbial communities in bats. PeerJ. 2019; 7: e6844–e6844. doi:10.7717/peerj.6844

10. McKenzie VJ, Song SJ, Delsuc F, Prest TL, Oliverio AM, Korpita TM, et al. The effects of captivity on the mammalian gut microbiome. Integr Comp Biol. 2017; 57(4): 690–704. doi:10.1093/icb/icx090

11. Carrillo-Araujo M, Tas N, Alcántara-Hernández RJ, Gaona O, Schondube JE, Medellín RA, et al. Phyllostomid bat microbiome composition is associated to host phylogeny and feeding strategies. Front Microbiol. 2015; 6: 447. doi:10.3389/fmicb.2015.00447

12. Marí Saéz A, Weiss S, Nowak K, Lapeyre V, Zimmermann F, Düx A, et al. Investigating the zoonotic origin of the West African Ebola epidemic. EMBO Mol Med. 2015; 7(1): 17–23. doi:10.15252/emmm.201404792

13. Illumina Inc. 16S Metagenomic Sequencing Library Preparation - Preparing 16S Ribosomal RNA Gene Amplicons for the Illumina MiSeq System [Internet]. 16S Metagenomic Sequencing Library Preparation Manual. 2013.

14. Klindworth A, Pruesse E, Schweer T, Peplies J, Quast C, Horn M, et al. Evaluation of general 16S ribosomal RNA gene PCR primers for classical and next-generation sequencing-based diversity studies. Nucleic Acids Res. 2013; 41(1). doi:10.1093/nar/gks808

15. Amir A, McDonald D, Navas-Molina JA, Kopylova E, Morton JT, Zech Xu Z, et al. Deblur Rapidly Resolves Single-Nucleotide Community Sequence Patterns. Gilbert JA, editor. mSystems. 2017; 2(2): e00191–16. doi:10.1128/mSystems.00191-16

16. Janssen S, McDonald D, Gonzalez A, Navas-Molina JA, Jiang L, Xu ZZ, et al. Phylogenetic Placement of Exact Amplicon Sequences Improves Associations with Clinical Information. mSystems. 2018; 3(3): e00021–18. doi:10.1128/mSystems.00021-18

17. Lozupone C, Knight R. UniFrac: a New Phylogenetic Method for Comparing Microbial Communities. Appl Environ Microbiol. 2005; 71(12): 8228. doi:10.1128/AEM.71.12.8228-8235.2005

18. Morton JT, Sanders J, Quinn RA, McDonald D, Gonzalez A, Vázquez-Baeza Y, et al. Balance Trees Reveal Microbial Niche Differentiation. Jansson JK, editor. mSystems. 2017; 2(1): e00162–16. doi:10.1128/mSystems.00162-16

19. Clayton JB, Vangay P, Huang H, Ward T, Hillmann BM, Al-Ghalith GA, et al. Captivity humanizes the primate microbiome. Proc Natl Acad Sci. 2016; 113(37): 10376 LP – 10381. doi:10.1073/pnas.1521835113

20. Kolodny O, Weinberg M, Reshef L, Harten L, Hefetz A, Gophna U, et al. Coordinated change at the colony level in fruit bat fur microbiomes through time. Nat Ecol Evol. 2019; 3(1): 116–24. doi:10.1038/s41559-018-0731-z

21. Phillips CD, Hanson J, Wilkinson JE, Koenig L, Rees E, Webala P, et al. Microbiome structural and functional interactions across host dietary niche space. Integr Comp Biol. 2017; 57(4): 743–55. doi:10.1093/icb/icx011

22. Kohl KD, Skopec MM, Dearing MD. Captivity results in disparate loss of gut microbial diversity in closely related hosts. Conserv Physiol. 2014; 2(1). doi:10.1093/conphys/cou009

23. Hughes GM, Leech J, Puechmaille SJ, Lopez J V., Teeling EC. Is there a link between aging and microbiome diversity in exceptional mammalian longevity? PeerJ. 2018; 6: e4174. doi:10.7717/peerj.4174

24. Lemieux-Labonté V, Tromas N, Shapiro BJ, Lapointe F-J. Environment and host species shape the skin microbiome of captive neotropical bats. PeerJ. 2016; e2430. doi:10.7717/peerj.2430

25. Nearing JT, Douglas GM, Comeau AM, Langille MGI. Denoising the Denoisers: an independent evaluation of microbiome sequence error-correction approaches. PeerJ. 2018; 6: e5364–e5364. doi:10.7717/peerj.5364

26. Degli Esposti M, Martinez Romero E. The functional microbiome of arthropods. PLoS One. 2017; 12(5): e0176573.

